# Denoising sparse microbial signals from single-cell sequencing of mammalian host tissues

**DOI:** 10.1101/2022.06.29.498176

**Authors:** Bassel Ghaddar, Martin J. Blaser, Subhajyoti De

## Abstract

We developed SAHMI, a computational resource to identify truly present microbial nucleic acids and filter contaminants and spurious false-positive taxonomic assignments from standard transcriptomic sequencing of mammalian tissues. In benchmark studies, SAHMI correctly identifies known microbial infections present in diverse tissues. The application of SAHMI to single-cell and spatial genomic data enables co-detection of somatic cells and microorganisms and joint analysis of host-microbiome ecosystems.

## Main text

The microbiome plays an integral role in healthy development, aging, and multiple diseases, however the nature of its influence in many contexts remains poorly understood^1^. This is because integrated analyses of host-microbiome ecosystems *in vivo* have been difficult to achieve in part due to the lack of relevant model systems^2^, technological barriers to directly profiling host-microbial interactions at high resolution in human tissues, and substantial heterogeneity across patients^3,4^.

While targeted detection of microbial antigens and 16s rRNA gene sequencing are standard microbiome profiling approaches, recent studies have reported detection of microbial nucleic acids in polyA-selected RNA sequencing of clinical samples from humans^5–7^. This is remarkable, but not surprising given increasing observations of polyadenylated transcripts in prokaryotes^8,9^ and that non-polyadenylated mammalian sequences are routinely captured in RNA-seq^10^. However, in samples that have low microbiome-biomass and are without matched and co-processed negative controls, denoising true microbial signals and removing contaminating species is a major challenge^11–13^. Important issues in the field that limit the use of emerging sequencing data to probe host-microbiome ecosystems include: (1) lack of gold standard contamination controls in most genomic experiments, (2) the usual lack of benchmarking for spurious, false-positive taxonomic assignments derived from human samples, and (3) the lack of benchmarking of metagenomic read capture from single-cell RNA sequencing (scRNA-seq) protocols.

To overcome these problems, we developed and benchmarked SAHMI (Single-cell Analysis of Host-Microbiome Interactions), a computational pipeline to identify microbial nucleic acid sequences and remove false positives and contaminants from bulk, single cell, and spatial transcriptomic data of mammalian tissues (**Fig. 1a**). SAHMI denoises and decontaminates the output of taxonomic classifiers (e.g. Kraken2Uniq^14,15^). It identifies taxa that are truly present in a tissue specimen by examining the relationship between the number of total and unique k-mers for each taxon, and it removes contaminants by comparing taxa profiles to an extensive negative control reference dataset. We now show that SAHMI successfully identifies known infections from scRNA-seq and spatial transcriptomic data sets from human tissues, and that microbes can be paired and jointly analyzed with somatic cells. SAHMI thus unlocks the potential for retrospective host-microbiome interaction analysis from a wealth of existing transcriptomic studies at different resolutions.

**Figure 1.**
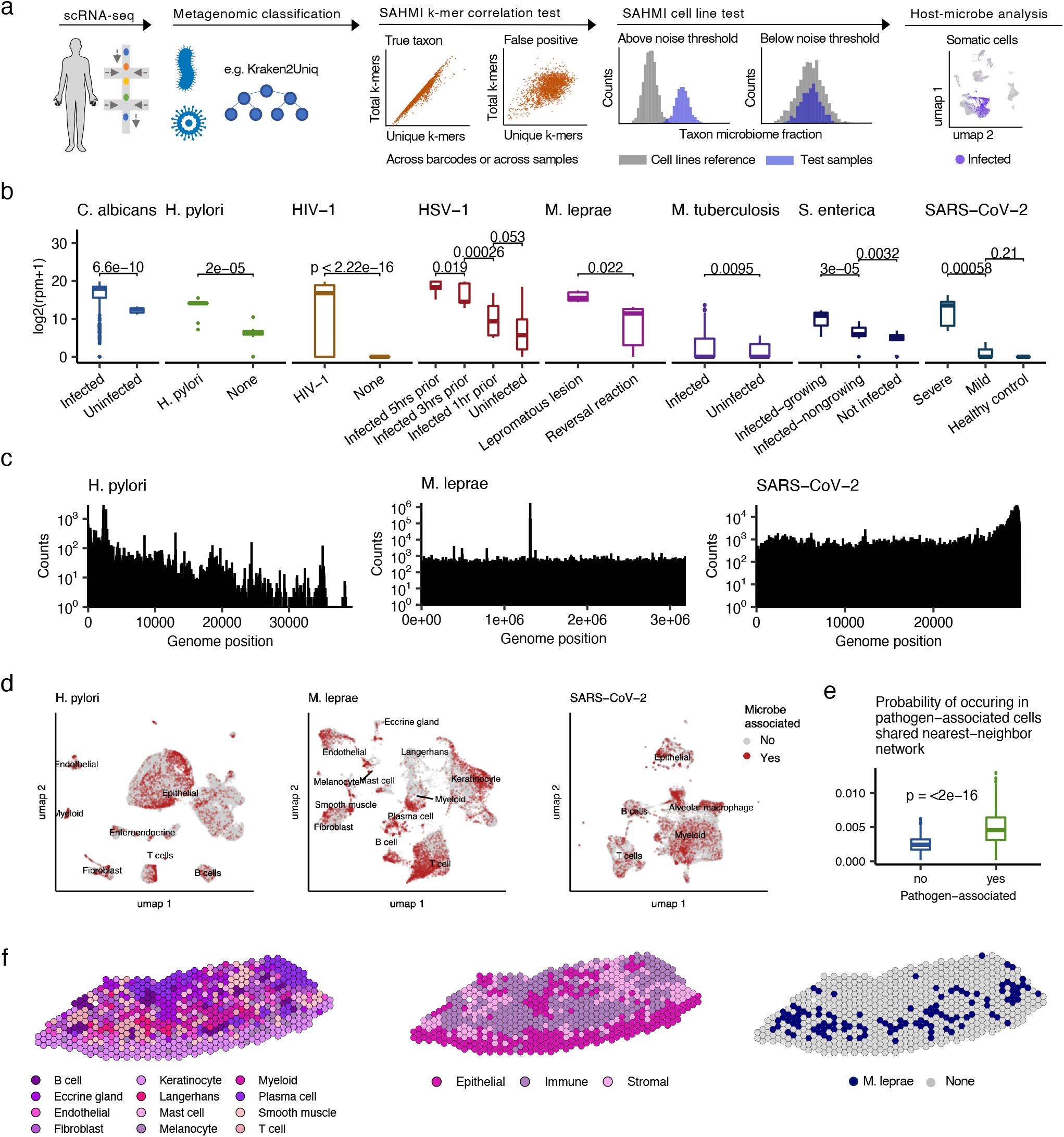
Host scRNA-seq captures true microbial reads. **a,** Schematic of the SAHMI pipeline. SAHMI identifies taxa that are truly present in tissues using a k-mer correlation test and identifies false positives and contaminants by comparing taxa distributions to an extensive negative control reference. **b,** Differential detection of known pathogens in infected samples in multiple studies. Boxplots show median (line), 25^th^ and 75^th^ percentiles (box) and 1.5xIQR (whiskers). Points represent outliers; all Wilcoxon testing. rpm, reads per million classified microbiome reads. **c,** Histograms of genomic mapping positions for the known pathogens in clinical studies. Reads map throughout the genomes. **d,** Uniform manifold approximation and projection (UMAP) plots of somatic cells from the clinical studies identifying cell types and cell barcodes paired with the known pathogen (red color). Microbe-associated host cells cluster together. Tissue sources: left, gastric biopsy; middle, skin; right: bronchoalveolar lavage fluid. **e,** Boxplots of the probability a pathogen-associated or unassociated host cell is in the same shared nearest-neighbor network as a pathogen-associated cell. Boxplots are as in (**b**). **f,** Spatial transcriptomic map of a lepromatous cutaneous lesion colored by predominant cell-type (left), cell compartment (middle), or presence of detected *M. leprae* (right).

First, we asked whether known pathogens could be differentially detected in a dosedependent manner in samples with known infection. We analyzed scRNA-seq data for human samples in which the host had a clinically verified infection^16–18^ or in which a pathogen was experimentally introduced^19–22^; these represented a variety of bacteria, fungi, viruses, and tissue types (**Table S1**). We used Kraken2Uniq^14,15^ to map all reads to a database of human and microbial genomes. The infection scRNA-seq data contained 10^7^-10^10^ reads, of which a mean of 1.3% (median 0.05%, standard deviation 3%) mapped to the microbiome. Across all studies, pathogen reads were successfully identified and were significantly increased in the samples with known or experimental infection with respect to the pathogen load (**Fig. 1b**). To validate that these assignments were not artifacts, we mapped pathogen reads from the clinical samples using STAR^23^, a slower but dedicated RNA-seq aligner. For all studies, >90% of Kraken2Uniq classified pathogen reads were aligned uniquely by STAR to regions throughout their respective genomes (**Fig. 1c**). These results indicate that true microbial nucleic acid sequences are quantitively captured in scRNA-seq.

Some scRNA-seq barcodes tagging microbial reads also tagged somatic cellular RNA, suggesting that these microbes and cells were co-localized. To determine if barcode sharing reflected the pairing of somatic and microbial cells *in vivo*, we examined the data for known microbe-cell-type specific interactions. Using the same samples from the clinical infection studies, we identified the somatic cell types and highlighted cells that were paired with a pathogen (**Fig. 1d**). Within a cell-type, and across all studies, microbe-associated cells generally clustered together, indicating shared, broad gene expression changes compared to unassociated cells. *Mycobacterium leprae* was most commonly found with T-cells, keratinocytes, and myeloid cells, consistent with its ability to directly infect macrophages and keratinocytes and reflecting the importance of T-cells in the immune responses related to granulomata^24^. *Helicobacter pylori* was mostly associated with a major subset of gastric epithelial cells and with mucosal immune cells. Severe acute respiratory syndrome related coronavirus 2 (SARS-CoV-2) was found broadly in epithelial and immune cells, and especially in alveolar macrophages (**Fig. 1d**). In all studies, pathogen-associated host cells were significantly more likely to be in the same shared nearest-neighbor network with other pathogen-associated cells compared to pathogen-unassociated cells, indicating their transcriptional similarity (Wilcoxon, p<2e-16; **Fig. 1e**).

Microenvironmental contexts of infection are important and can be studies with SAHMI by utilizing shared barcodes in spatial transcriptomic data. We analyzed spatial sequencing data of a lepromatic lesion and identified the predominant cell-type from each spatial spot as well as spots that captured *M. leprae* RNA (**Fig. 1f**). Inspection of the spatial map shows *M. leprae* primarily overlapping with immune regions of the tissue, reflecting granuloma response to infection. These data demonstrate how the increased precision afforded by molecular barcoding localizes microbes to specific host cells and can enable downstream examination of cell-type specific gene expression as it relates to the presence of a microorganism.

While these analyses showcase the reliable detection of known microbes using SAHMI, filtering contaminants and spurious false positive taxonomic assignments is crucial for the study of tissues with unknown microbial burden. In above analyses, a mean of 4735 other species (range 801-7084) were classified in the benchmark datasets, and this number correlated with the total number of reads per study (r=0.7, p=0.08, **Fig. 2a**). These included common contaminants such as *Mycoplasma* spp. and *Cutibacterium acnes*, but also unexpectedly ubiquitous species such as *Xanthomonas euvesicatoria* and proteus virus Isfahan. This was especially surprising because five of the eight benchmark studies had pathogens experimentally introduced under strict aseptic technique and suggests that the majority of reported taxa were spurious false-positive assignments. We also observed that reported microbial profiles differed significantly depending on the mapping parameters used. For example, mapping reads from the clinical studies to the microbiome alone without including the host genome led to significantly increased reads that mapped to bacteria in general but to only negligible differences in the number of reads that mapped to the verified pathogen (t-test p<1e-4, **Fig. 2b**) – again underscoring the presence of false-positives as well as the importance of including all relevant reference genomes during taxonomic classification. The number of unique k-mer sequences assigned to a taxon has been used to filter false positives; however, we observed a wide range in these values across taxa, and they did not clearly distinguish the known pathogens (**Fig. 2c**).

**Figure 2.**
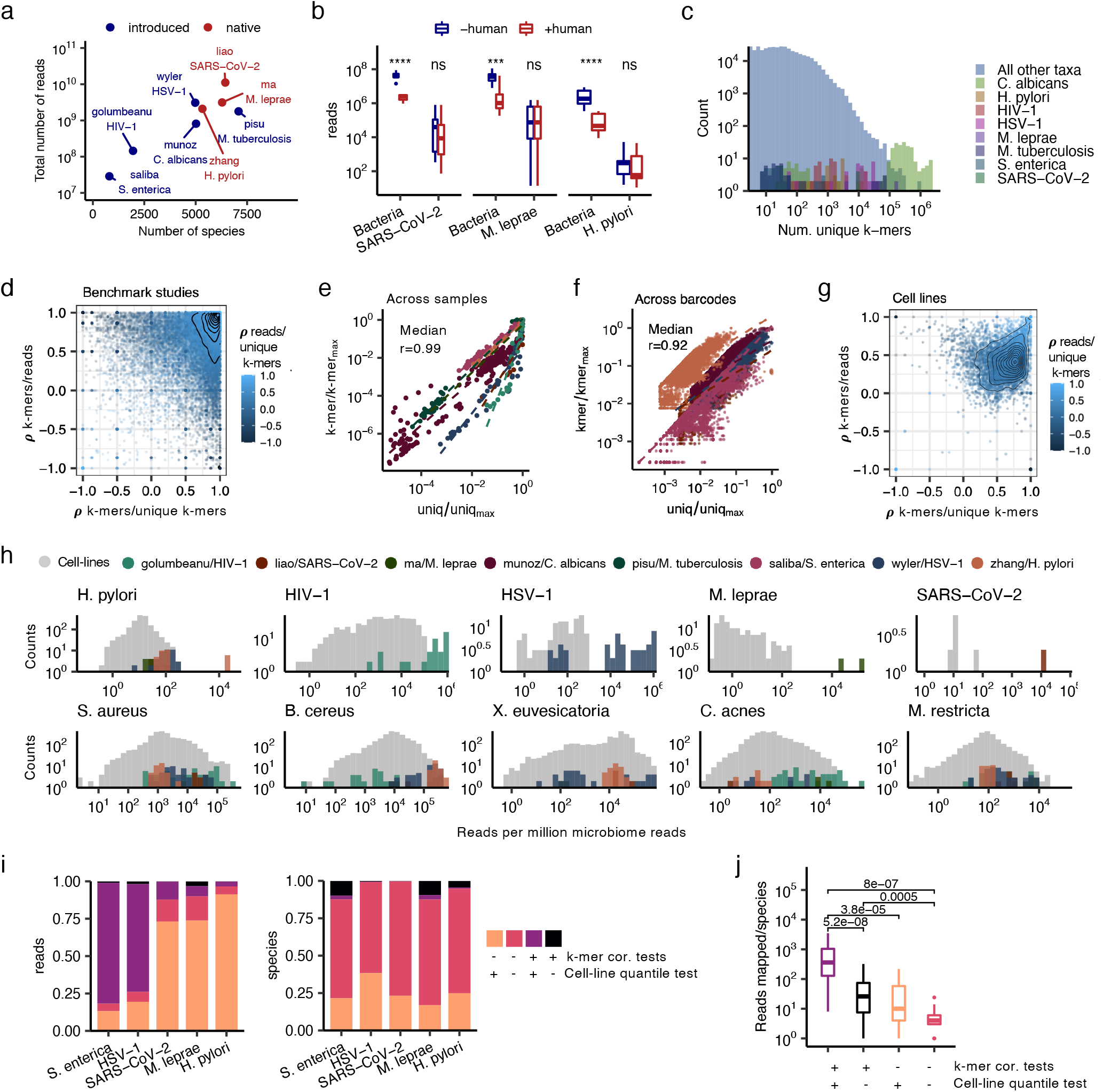
Distinguishing true signal from false positives and contaminants. **a,** Scatter plot showing the total number of sequencing reads and species detected in each study. Blue, experimentally introduced pathogen; red, existing infection in a human tissue. **b,** Boxplots showing significantly increased reads assigned to bacteria when the human genome is not included as a reference during taxonomic classification. Boxplots show median (line), 25^th^ and 75^th^ percentiles (box) and 1.5xIQR (whiskers). Points represent outliers; t-tests; **** p<1e-3; *** p<1e-2; ns, not significant. **c,** Histogram of the number of unique k-mers per sample assigned to the known pathogens and to all detected species in the benchmark studies. **d,** Scatter plot of the k-mer correlation tests for species in the benchmark studies. Each point represents an individual species. Correlations are run across samples within a study. The x-axis represents the Spearman correlation values between #k-mers vs. #unique k-mers. The y-axis represents the correlation value between #k-mers vs. #reads; colors represent correlation value between #reads vs. #unique k-mers. Lines represent contour density. **e,** Scatter plot of the number of total and unique k-mers for the known pathogens. Each point represents a sample. The points are colored by study as detailed below the plot. **f,** Scatter plot of the number of total and unique k-mers for the known pathogens. Each point represents an individual barcode. Points are colored as in (**e**). **g,** Similar to (**d**) but for taxa detected in cell line experiments. **h,** Overlaid histograms of reads per million microbiome reads assigned to the known pathogens (top row) and to example contaminants (bottom row) in the benchmark studies as well as in the cell line data. Bars are colored by study of origin. **i,** Bar plots of relative proportions of species (left panel) and reads (right panel) that passed or failed the k-mer correlation and cell-line quantile tests. **j,** Boxplots showing the number of reads per taxon mapped by STAR for microbial reads from the leprosy study grouped by k-mer correlation and cell line quantile test results. Colors are as in (**i**) and boxplots are as in (**b**). Wilcoxon testing performed.

The issue of false positive taxonomic assignments is well-known^11,25^; these may arise from multiple sources, including sequence homology, sequencing errors, off-target amplification, or mapping errors. To limit such identifications, we posited that in the setting of low-microbiome biomass, a proportional number of total and unique RNA transcripts will be captured for true species. When we examined the pairwise correlations across samples between the numbers of reads, k-mers, and unique k-mers assigned to each species in each benchmark study, we found a wide spectrum of values (**Fig. 2d**), with the true pathogens having extremely high correlation values (median r>0.92, p<2e-16, **Fig. 2e, Table S1**). The strong correlation between the number of total and unique k-mers also held true for the known pathogens across barcodes, allowing us to identify microbes in individual samples (**Fig. 2f**). These k-mer correlation tests served as a significant filter; while 6063 species were classified in the tested samples, only 1207 species had significant values in all tests.

We next sought to identify contaminants based on the observation that contaminants appear in higher frequencies in negative control samples^12,13^. In the absence of matched controls, we posited that sterile cell line data could serve as a substitute. We profiled the microbiome from publicly available RNA-seq data for 2,491 samples involving >1500 human cell lines representing healthy and diseased tissues from >400 sources from around the world (**Table S2**), thereby creating a negative control resource and identifying common contaminants and false positives (**Table S3**). From these studies, we identified a mean of 1035 species per sample (range: 124-6731). Our k-mer correlation tests again found a range of values for the cell line taxa (**Fig. 2g**). These correlations were significantly weaker than those from the benchmark studies (Wilcoxon, p<2e-16) due to the cell line microbiome data being enriched in false positives. The most ubiquitous species included cutaneous microbiota and common environmental or laboratory species (**Table S3**).

Comparing taxa reads counts from the benchmark studies to their distribution in the cell line data using a quantile test clearly distinguished true positive signal and identified background contamination or noise in all studies (Wilcoxon p<2e-16, **Fig. 2h**). The known pathogens had significantly higher reads per million microbiome reads compared to what was found in the cell line data, whereas most taxa did not. Our procedures significantly reduced false positives: only a minority of all reported species (median: 2.8%, range: 0.26-22%) and initially classified microbial reads (median: 12%, range: 3.2-81%) passed both the k-mer correlation and cell line quantile tests (**Fig. 2i**). In the studies with an introduced pathogen (e.g. HSV-1, *S. enterica*), only a median of 3 species per sample passed our pipeline (range: 1-6). To validate that our pipeline enriched for true signal, we used STAR^23^ to map reads for a subset of species from the skin leprosy study and found that species that passed both k-mer correlation and cell line quantile tests had significantly more mapped reads than all other initially reported taxa (**Fig. 2j**), despite comprising a minority of reads and species. These validation and benchmarking analyses collectively demonstrate how SAHMI can systematically enrich for true taxa and eliminate contaminants and false positives across a range of datasets from diverse tissues.

SAHMI offers a resource to study microbial-cell-type-specific interactions at single cell resolution and in spatial contexts *in vivo*. It enables the detection, localization, and association of microbes with host cell gene expression in existing and new human or other mammalian scRNA-seq data from a variety of tissue types, including cancer^30^. SAHMI is available on Github (https://github.com/sjdlabgroup/SAHMI).

## Methods

### SAHMI pipeline for microbiome detection and denoising from transcriptomic data

We developed SAHMI, a statistical pipeline for detection of microbes and analysis of host-microbiome interactions from scRNA-seq and other transcriptomic data. After sequencing and taxonomic classifications, SAHMI has two primary functions: (1) it identifies true microbial signal by running correlation analyses across barcodes and samples (k-mer correlation tests), and (2) it filters contaminants and false positives by comparing metagenomic counts to distributions of profiles in negative control samples (cell line quantile test). This enables systematic retrospective identification of microbes in host tissues. Downstream analysis of associations can be done at the sample level or at the level of individual cells for somatic cells and microbes that are tagged with the same cell barcodes.

#### Taxonomic classification

metagenomic classification of paired-end reads from bulk RNAseq, scRNA-seq, or spatial transcriptomic sequencing fastq files can be performed using a k-mer based mapper that identifies a taxonomic ID for each k-mer and read. While SAHMI can work with any k-mer-mapper, we reported the results for SAHMI with Kraken2Uniq^14,15^, a popular and benchmarked tool which finds exact matches of candidate 35-mer genomic substrings to the lowest common ancestor of genomes in a reference metagenomic database. It is essential that all realistically possible genomes are included as mapping references at this stage, or that host mappable reads are excluded. The required outputs from this step are: a Kraken summary report with sample level metagenomic counts, a Kraken output file with read and k-mer level taxonomic classifications, and raw sequencing fastq files with taxonomic classification for each read, or the equivalent data.

#### Barcode level signal denoising (barcode k-mer correlation test)

SAHMI first extracts microbiome reads from the raw data given their taxonomic IDs and removes reads that contain k-mers that map to the host genome. Next, for each taxon in the sample, SAHMI identifies the corresponding reads and removes reads with less than a default of 50% of the k-mers mapped directly to the taxon or to a parent taxon in its lineage. While analyzing scRNA-seq data, the cell barcode ID is used to identify reads originating from the same droplet. The number of total and unique k-mers mapping to the taxon or its lineage is then tabulated. For computational efficiency, a default of 1000 barcodes per taxon are randomly sampled. The Spearman correlation between the number of total and unique k-mers across barcodes for each taxon is computed. SAHMI reports the correlation and p-value and recommends removing taxa with non-significant correlations. This enables identification of true microbes in an individual sample.

#### Sample-level signal denoising (sample k-mer correlation test)

the correlation analysis is also conducted across samples when possible. The Kraken report tabulates the total number of reads, minimizers (k-mers), and an estimate of unique k-mer counts for each taxon, or the equivalent data can be obtained from mappers. For each taxon, SAHMI correlates the number of k-mers vs. unique k-mers, reads vs. k-mers, and reads vs. unique k-mers across all samples in a study. True taxa are identified as those having significant positive Spearman correlation values and p-values for all three tests.

#### Identifying contaminants and false positives (cell line quantile test)

these can be identified in the SAHMI workflow based on the widely observed pattern that contaminants appear at higher frequencies in low concentration or negative control samples^12,13^. We observed that this pattern also extends to false positive assignments. In the absence of experimentally matched negative controls, we provide a negative control resource comprised of microbiome profiles from 2,491 sterile cell experiments from around the world. For each taxon in a test sample, SAHMI compares the fraction of microbiome reads assigned to the taxon [i.e. taxon counts/sum(all bacterial, fungal, viral counts), in reads per million] to the microbiome fraction assigned to the taxon in all cell line experiments. Using the microbiome fraction comparison normalizes for experiments having a varying number of total sequencing reads or varying underlying contamination. SAHMI tests whether the taxon microbial fraction in the test sample is > 99^th^ percentile (by default) of the taxon’s microbiome fraction distribution in cell line data using a one-sample quantile test. Taxa whose counts fall within the cell line distribution are identified as below the cell-line noise threshold. Users may choose how stringently to select the quantile threshold for significance testing.

#### Quantitation of microbes and creating the barcode-metagenome counts matrix

after identifying true taxa, reads assigned to those taxa are extracted and passed though a series of filters. ShortRead is used to remove low complexity reads (< 20 non-sequentially repeated nucleotides), low quality reads (PHRED score < 20), and PCR duplicates tagged with the same unique molecular identifier and cellular barcode. Non-sparse cellular barcodes can be selected by using an elbowplot of barcode rank vs. total reads, smoothed with a moving average of 25, and using a cutoff at a change in slope < 10^-3^, in a manner analogous to how cellular barcodes are typically selected in single-cell sequencing data (CellRanger (10x Genomics), Drop-seq Core Computational Protocol v2.0.0 (McCarroll laboratory)). Lastly, the full taxonomic classification of all resulting reads and the number of reads assigned to each clade are tabulated.

### Assembling the negative control cell lines microbiome data

The Sequence Read Archive (SRA) was queried using the following search: ((((“public”[Access]) AND “rna seq”[Strategy]) AND “transcriptomic”[Source]) AND cell line) AND “Homo sapiens”[orgn: txid9606] to identify sequencing runs with human cell lines. This resulted in 52,397 sample entries. We then selected for samples with “library selection = cDNA OR library selection = PolyA”, and we removed experiments with mouse strain information, experiments involving infection, runs without a submitting center name, and runs with “cell line” designated as “none”. From each remaining submitting center, we randomly selected 5 runs. We combined these run IDs with the run IDs for the complete Cancer Cell Line Encyclopedia^26^. We downloaded raw RNA-seq fastq files for these samples and profiled their microbiome using Kraken2Uniq^14,15^. Samples were retained in the data base only if >90% of the reads were classified as human (taxid=9606). We then manually checked each sample’s metadata to ensure they did not involve infection or stimulation with microbial agents. The resulting samples and their metadata are tabulated in **Table S2** and cell line microbiome data are available in **Table S3**.

### True positive datasets selection, metagenomic mapping, and scRNA-seq data processing

We analyzed scRNA-seq data from patient samples with the following clinically verified infections: *Mycobacterium leprae* (skin)^17^, *Helicobacter pylori* (stomach)^18^, and severe acute respiratory syndrome coronavirus 2 (SARS-CoV-2, bronchoalveolar lavage fluid)^16^, in addition to data from scRNA-seq experiments for tissues in which the following pathogens introduced: *Candida albicans* (PBMCs)^20^, *Salmonella enterica* (PBMCs)^19^, *Mycobacterium tuberculosis* (lung)^21^, human alpha herpesvirus 1 (HSV-1) (PBMCs)^27^, and human immunodeficiency virus 1 (HIV-1) (PBMCs)^22^. Taxonomic classification was done using Kraken2Uniq^14,15^ using a combined reference database including the human genome and all bacterial, fungal, and viral reference genomes recorded in RefSeq as of April 2022. Microbial read alignment was done for a subset of samples using the STAR^23^ RNA-seq aligner with the following parameters: alignIntronMax=1 and outFilterScoreMinOverLread=0.05. For the clinical datasets, scRNA-seq data were processed using the standard Seurat^28^ pipeline. Cell types were identified by comparing cluster marker genes to the PanglaoDB^29^ reference database. Cell-types in the spatial transcriptomic map of a lepromatic lesion were identified as follows. Marker gene for each cell-type were identified from scRNA-seq data of lepromatic lesions using the FindAllMarkers function with logfc.threshold=1 and min.pct.diff=0.25. Cell-type scores for each spatial spot were computed as the mean expression of cell-type marker genes, and these were scaled per cell-type. The predominant cell-type at a spatial spot was identified as the cell-type with the highest score at that spot.

### Statistical analyses

All statistical analyses were performed using R version 3.6.1. All p-values were corrected for false-discovery rate (fdr) for multiple hypotheses using the p.adjust function with method= “fdr”, unless otherwise stated. The ggpubr package (https://github.com/kassambara/ggpubr) was used to compare group means with nonparametric tests and to perform multiple hypothesis correction for statistics that are noted in the figures. P-values reported as <2.2×10^-16^ result from reaching the calculation limit for the native R statistical test functions and indicate values below this number, not a range of values. Barcode level analyses (**Fig. 2g, 2j**) were done for the benchmark studies in which reads had identifiable cell barcodes, unique molecular identifiers, and poly-A sequences. Comparisons to the human cell line negative control dataset were done for the benchmark studies that used human cells.

## Supplementary Materials

**Table S1.** Summary of benchmark studies analyzed.

**Table S2.** Summary of cell line experiments analyzed.

**Table S3.** Cell lines microbiome reference

## Acknowledgments

We acknowledge the Office of Advanced Research Computing (OARC) at Rutgers, The State University of New Jersey for providing access to the Amarel cluster URL: https://it.rutgers.edu/oarc.

## Funding

National Institutes of Health grant R21CA248122 (SD)

National Institutes of Health grant U01 AI22285 (MJB) Sergei Zlinkoff Foundation (MJB)

Canadian Institute for Advanced Research (MJB)

National Institutes of Health, National Center for Advancing Translational Sciences, Rutgers Clinical and Translational Science Award TL1TR003019 (BG)

## Author contributions

BG and SD conceived the study. BG designed and performed all data analyses. BG, MJB, SD interpreted the data and wrote and revised the manuscript.

## Data and materials availability

The cell lines microbiome negative control dataset is available in **Supplementary Table S3**. The SAHMI pipeline is available on our Github: https://github.com/sjdlabgroup/SAHMI.

